# First Report of *Lactoria cornuta* from Karnataka, West coast of India–Role of tropical cyclones in Range Extension

**DOI:** 10.1101/2021.06.28.450260

**Authors:** A.N. Muhammed Zafar Iqbal, Gurudev Mali, Raghavendra Dollin

## Abstract

India is known for its rich biodiversity and is fortunate to have several endemic species from different classes of vertebrates. India is home to 7.5% of the global fish diversity, with 91 endemic species of ray finned fishes, the actinopterygians. Some fish species have never been reported until recently, one such example is the long-horned Cowfish (Ostracidae), best known for inhabiting only certain areas of the Indo-Pacific oceans. However, this fish has been reported from Bay of Bengal, Costs of Goa and Lakshadweep. This is the first time this fish has been found on the Coasts of Karnataka, west coast of India. Given its morphology, the migration seems highly improbable since it can only endure lethargic swimming. In this article, we have explored the role of other external forces that could have contributed to its range extension in the east and west coast of India. In this paper we are exploring the role of Super Cyclones as a vital force in determining the expansion of its range. The fact of collection longhorn cowfish for the first time from the coast of Karwar coincides with the passage of Cyclone Ockhi on the west coast of India this suggests the possible role of super cyclone in its range extension. A year later, another specimen was recovered from a location very close to the first, which indicates the successful establishment of *Lacturia cornuta* in its new environment. Related morphometric and meristic studies of our specimens are consistent with previous studies reported in the Bay of Bengal.

## Introduction

India has 7.5% of the world’s fisheries diversity with 2,443 species from 927 genera (230 families of 40 orders). The Andaman Archipelago and the Nicobar Archipelago are the most numerous (1431 species), followed by the east coast of India (1121 species) and the west coast (1071 species) (Van der Elst, 1993; Chaudhuri, 2004; Berra, 2001; Leveque. et. al. 2008; Mishra and Gopi, 2015). The fishing industry makes an important contribution to the social and economic status of people living in coastal areas. In addition, India has 91 endemic fish species, the majority of which are found on the east coast of India. Unfortunately, an almost equal number of species have been introduced into the IUCN Near-Threatened category. The threat to fish diversity is primarily due to habitat degradation and non-scientific harvesting of fish resources.

Fish have a very large number of poisonous or venomous species, indeed more than 50% of venomous and venomous vertebrates are actually fish (Smith and Wheeler, 2006; John et al. 2014). Consequently, the number of venomous and poisonous fish are divided into several orders of ray-finned fish (Church and Hodgson, 2002; Halstead, 1970; 1988; Haddad et al. 2003; Haddad Jr V, Martins, 2006; Haddad Jr V et al. 2009; Vetrano et al 2002). Almost all tetradontiform fish, which constitute approximately 5% of total fish, are poisonous or toxic. Some of the familiar representatives found off the coast of India are Porcupinefishes (Diodontidae), Puffers (Tetradontidae), spikefishes (Triacanthodidae), triple spine fishes (Triacanthidae), triggerfishes (Balistadae), the boxfishes (Ostracionidae) and molas (Molidae). Many have powerful toxins in any of the organs such as the hepatopancreas, ovary or intestine, but certain species secrete toxins from their dermal glands. Tetrodotoxin is one of the most potent neurotoxins affecting predominantly neuromuscular transmission and sympathetic nervous system (Gopalakrishnakone and Malhotra, 2017; Smith and Wheeler, 2006). Beyond the associated risk, these fish are unpleasant, preventing them from being exploited commercially, whether for food or other commercial purposes.

The Ostraciidae family is one of ten families of this order with 33 species (12 existing genera) in the world. The Ostraciidae fishes have a peculiar square, box-shaped body and are very similar to pufferfishes and filefishes. The mucus contains a potent toxin, ostracitoxin (Pahutoxin), secreted from club cells in the skin epidermis (Halstead, 1970; 1988). Along the coastlines of India, nine species from four genera have been documented (Matsuura 1998; 2015). The Longhorn cowfish (*Lactoria cornuta cornuta*) is extensively distributed along the coasts of the Red Sea, Japan, South Korea and Australia at a depth of 1 to 50 meters.This species prefers habitats such as coral reefs, reef flats, and muddy or sandy coastlines (Lieske and Myers, 1994; Ambak et al.2010; Allen and Erdmann, 2012; Hutchins 1984). Porcupine and pufferfish of the Ostraciidae family continue to be reported on the east and west coasts of India. Nevertheless, the longhorn cowfishes were reported only from selected pockets of the Bay of Bengal and also from two locations along the west coast of India. The first record of Cowfishes emerged from the Lakshdweep islands in the Arabian sea followed by coast of Goa in 2013 (Vijay Anand & Pillai 2003: Hegde et al. 2013). During same period it was also reported for the first time from Bangladesh (2013) and later in 2017 it was reported from Odisha and Pondicherry. (Both reports from the Bay of Bengal) (Khan, 2013; Behera, 2017). In this article, we report for the first time the collection of Longhorn Cowfishes from Karnataka, west coast of India. This article also examines the factors, which might have contributed to the range extension of this species.

## Materials and Method

### Collection of first Specimen

The first specimen of Longhorn cowfish, *Lactoria cornuta cornuta* was caught (Fig. 2.0) alive from a small water puddle at Ravindranath Tagore beach, Karwar (14.809863 N and 74.125204 E) between 5 PM to 5:30 PM on 30 March 2018 during routine field studies. Collected specimen identified as longhorn cowfish, necessary photography and videography was done in the field itself to capture its activities. Live specimen was carefully collected and transported to the laboratory in plastic container with 15 liters of seawater for further studies. Notwithstanding to all our possible efforts, the specimen could be maintained alive for more than one hour in the laboratory condition. The specimen designated as **WCI18**, which stands for **W**est **C**oast of **I**ndia 20**18**.

**Fig 1.0.**
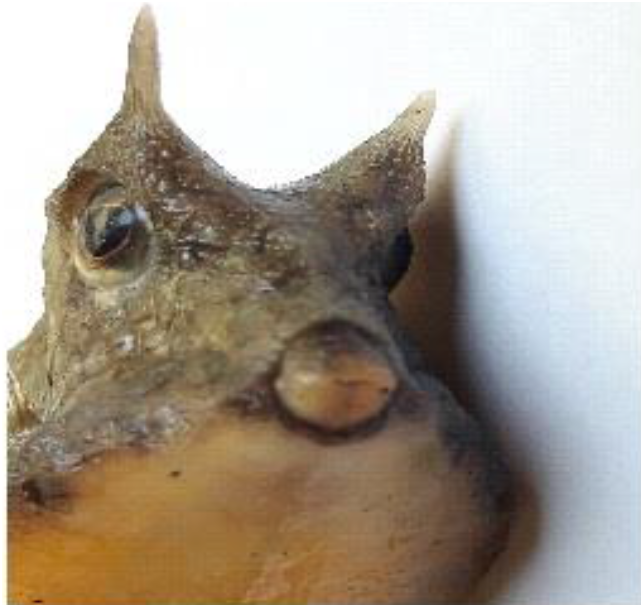
Sub terminal Mouth - Cowfish!!!

**Fig 2.0.**
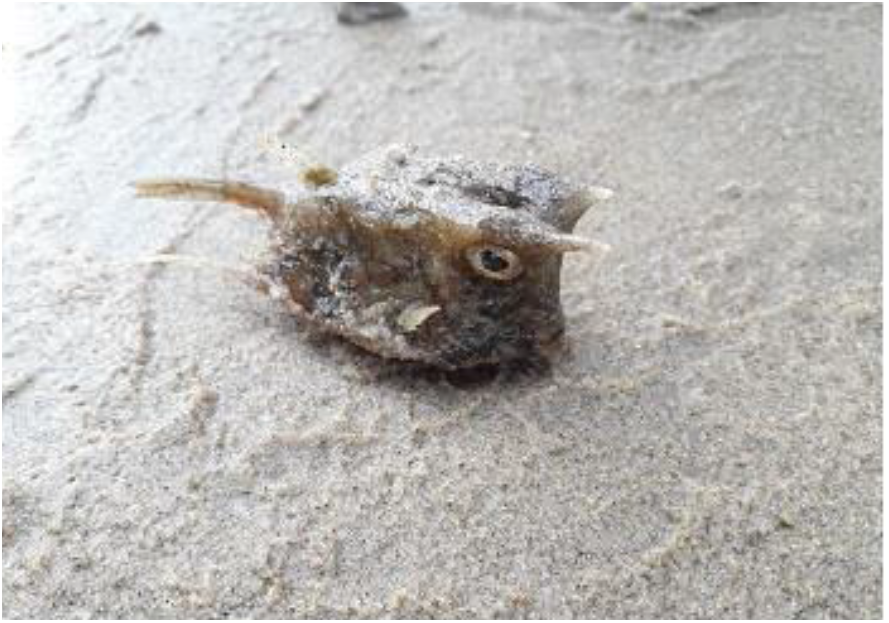
Cowfish - Collection Site.

**Fig 3.0.**
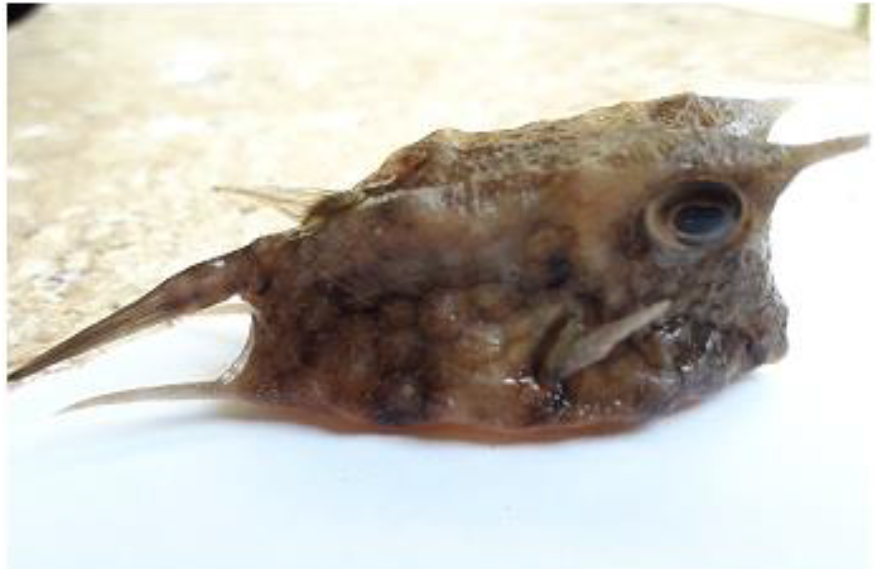
Lateral View.

**Fig 4.0.**
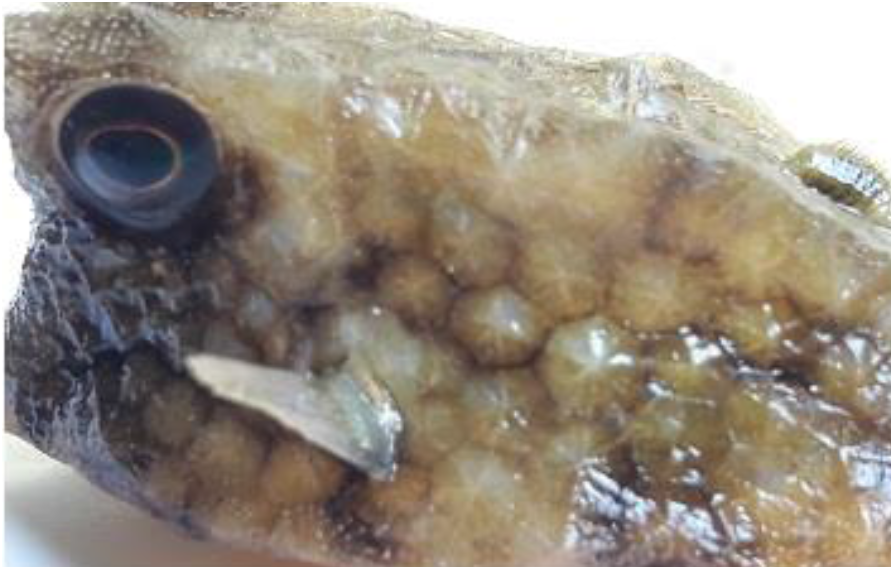
Plate like Dermal Scales.

**Fig 5.0.**
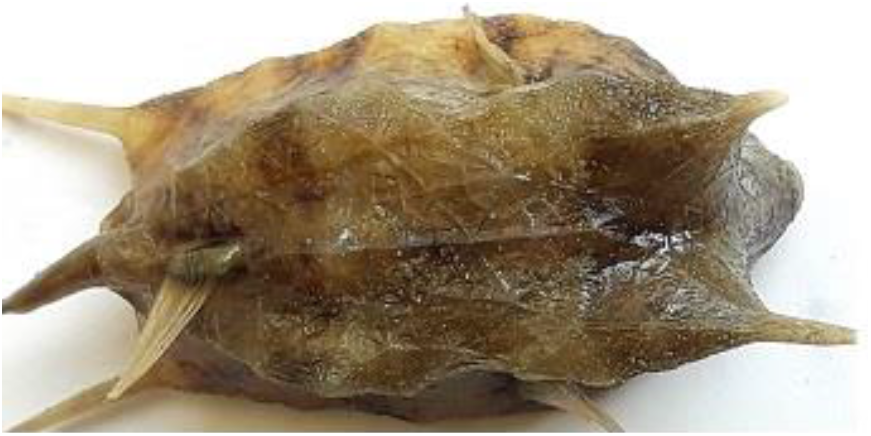
Dorsal View.

**Fig 6.0.**
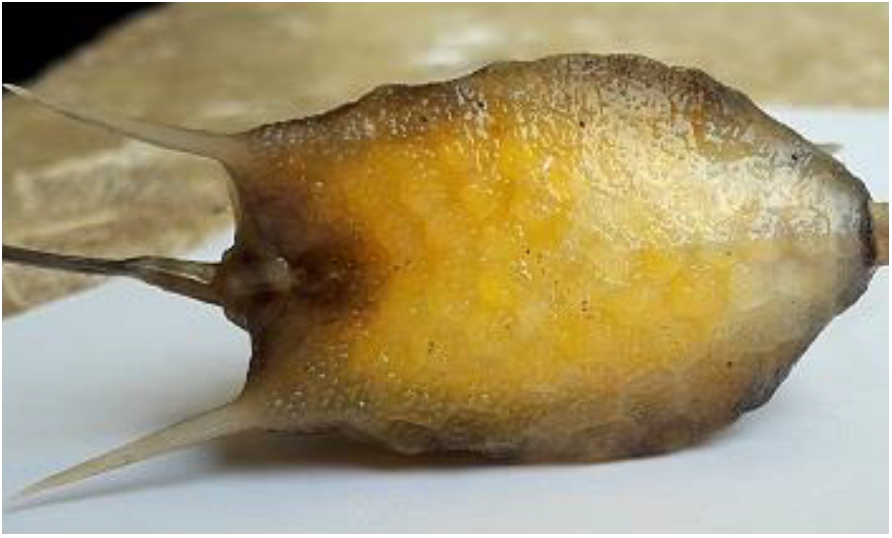
Lateral View.

**Fig 7.0.**
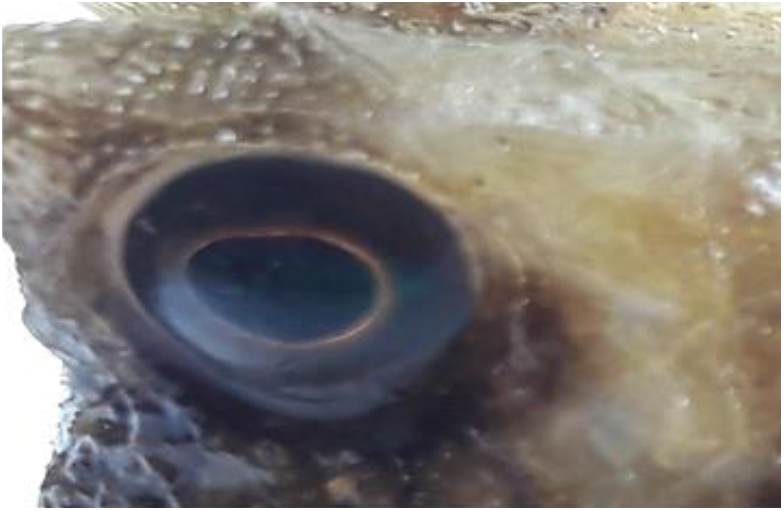
Orbit.

**Fig 8.0.**
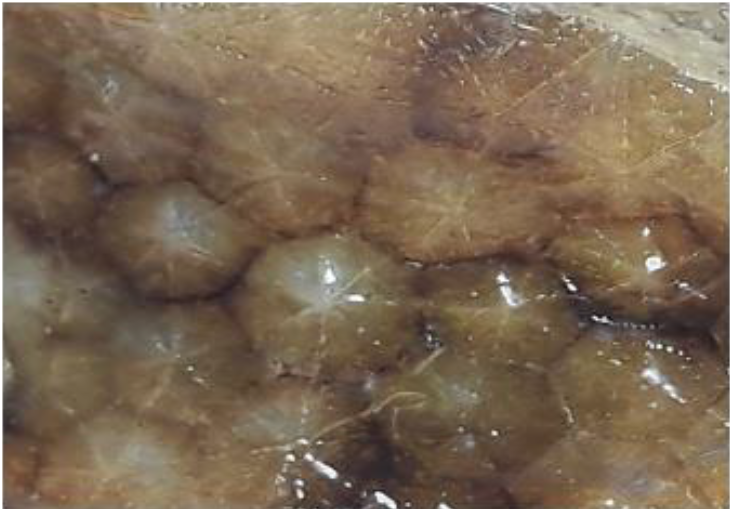
Plate like Dermal Scales.

### Morphometric and Meristic studies

The specimen subjected to morphometric and meristic characterization by adopting the procedure given by Hutchins and Randall (1982). Necessary morphometric measurements recorded by using digital caliper and meristic studies done by recording number of rays in the paired and median fins Identification. Both morphometric and meristic data tabulated systematically.

### Collection of second Specimen

After the collection of first specimen (WCI18), we were vigilant and continuously in search of more number of species. Fortunately, second dead but intact specimen was collected at around 7 AM on 18th February 2019 from Majali beach (14.88126 N & 74.10631 E) 10 km south of Karwar town. The specimen, transported to the laboratory and designated as **WCI19** (**W**est **C**oast of **I**ndia 20**19)** and necessary Photographs were taken. Morphometric and meristic studies carried out by following the Hutchins and Randall (1982) as described for WCI18 specimen and the data recorded and tabulated.

## RESULTS

The specimens conclusively confirmed as longhorn cowfish *Lactoria cornuta* by using identification keys provided by various authors (Hutchins and Matsuura, 1984; Randall et al.,1990; Allen, 2012; Ambak 2010).

### First specimen WCI18

The specimen WCI18 collected from Ravindranath Tagore beach, studied in detail and all morphometric and meristic characters measured and tabulated (Table 1.0.) Important morphometric measurements such as body length, body depth, eye diameter & inter-orbital length, fin lengths were recorded. During it was found that anterior dorso-lateral horns were unequal in length (Right measured 1.5 cm and left 0.6 cm) (Fig.11.) However, posterior dorso-lateral horns were equal both measuring 2.4 cm. (Fig. 12.) The right and left anterior horns spaced closely with gap of 2.3 cm while the posterior horns were slightly distantly spaced with 3.4 cm. gap in between. All rays of median fins (Dorsal, anal and caudal) (Fig. 10) and paired (Pectoral fin Fig. 9.0) counted carefully and meristic characters depicted in table 2.0.

**Table 1.** Morphometric characters of *Lactoria cornuta* specimens (WCI18 & WCI19)

| Sl.No. | Character | Measurements in cm |  |
| --- | --- | --- | --- |
|  |  | WCI18 | WCI19 |
| 1 | Total length (TL) | 12.6 | 19.2 |
| 2 | Standard length (SL) | 7.9 | 12.8 |
| 3 | Head length (LC) | 2 | 3.3 |
| 4 | Pre Orbital Distance (PRO) | 0.4 | 0.8 |
| 5 | Maximum body depth (Hx) | 3.8 | 3.8 |
| 6 | Minimum body depth (Hm) | 2.3 | 2.3 |
| 7 | Caudal peduncle length (LPC) | 2.8 | 2.8 |
| 8 | Eye diameter (OH) | 0.9 | 1.2 |
| 9 | Inter-orbital Distance (IO) | 1.1 | 3.1 |
| 10 | Pre-dorsal Distance (PD) | 5.3 | 7.6 |
| 11 | Pre-anal length (PA) | 5.8 | 9.2 |
| 12 | Pre-pectoral Distance (PPEC) | 2.3 | 3.6 |
| 13 | Depth of Dorsal fin (DDF) | 1.5 | 2.2 |
| 14 | Depth of Anal fin (DAF) | 1.3 | 2 |
| 15 | Length of Pectoral fin (LPF) | 1.6 | 2.7 |
| 16 | Length of Caudal fin (LCF) | 6.5 | 7.1 |
| 17 | Pectoral fin base length 1 | - | 1 |
| 18 | Pectoral fin base length 2 | - | 1 |
| 19 | Caudal fin base length | - | 1.4 |
| 20 | Width of caudal fin | - | 7.7 |
| 21 | Dorsal fin base length | - | 1.2 |
| 22 | Dorsolateral Anterior Horn | 1.5 R & 0.6 L | 2.0 L/R |
| 23 | Ventrolateral Posterior Horn | 2.4 R/L | 2.0 L/R |
| 25 | Distance between two horns at head region | 2.3 | 2.7 |
| 26 | Distance between two horns at anal region | 3.4 | 3.6 |

**Table 2.** Meristic characters of *Lactoria cornuta* specimens (WCI18 & WCI19)

| Sl.No. | Meristic Characters | Specimen |  |
| --- | --- | --- | --- |
|  |  | WCI18 | WCI19 |
| 1 | Dorsal spines | 0 | 0 |
| 2 | Dorsal fin rays | 9 | 9 |
| 3 | Anal spines | 0 | 0 |
| 4 | Anal fin rays | 9 | 9 |
| 5 | Caudal fin rays | 10 | 10 |
| 6 | Pectoral fin rays | 10 | 10 |
| 7 | Weight | Not recorded | 81.89 gm |

**Fig 9.0.**
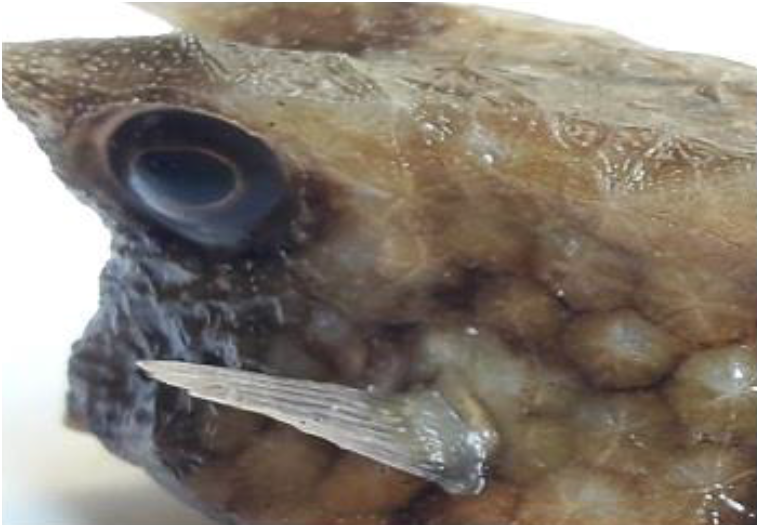
Pectoral Fin.

**Fig 10.0.**
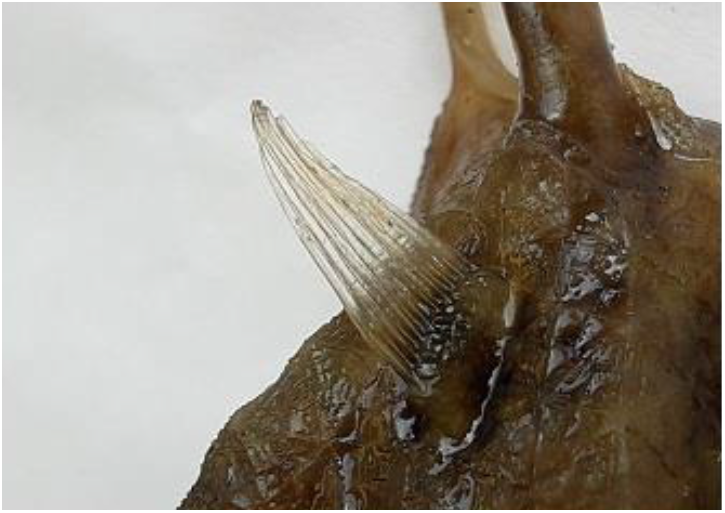
Dorsal Fin.

**Fig 11.0.**
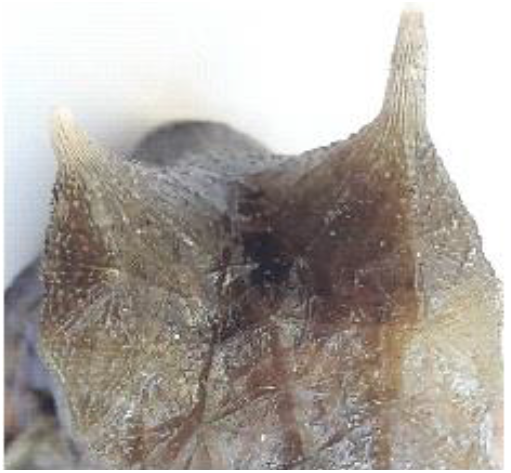
Anterior Horn.

**Fig 12.0.**
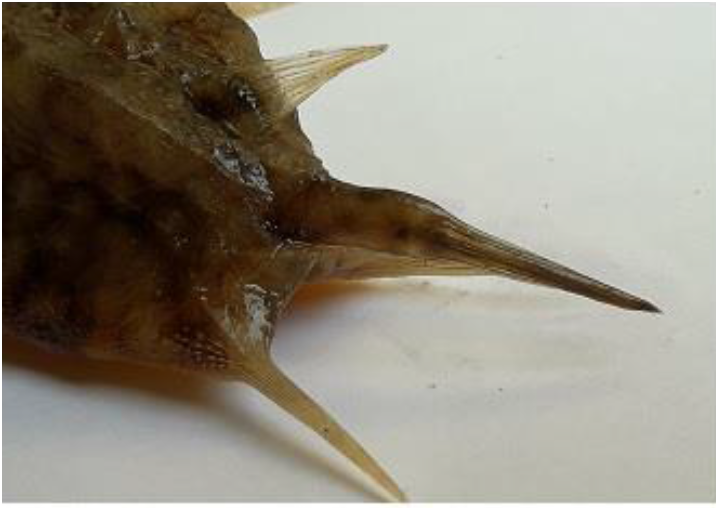
Posterior Hom.

**Fig 13.0.**
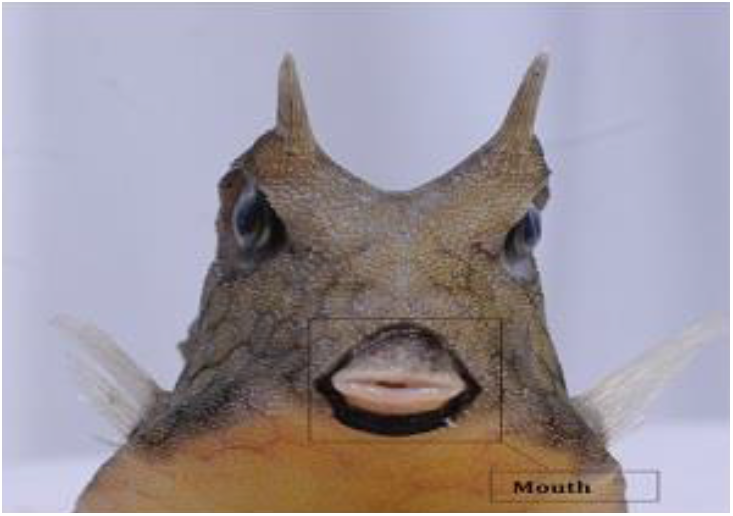
Front View - Cowfish!!

**Fig 14.0.**
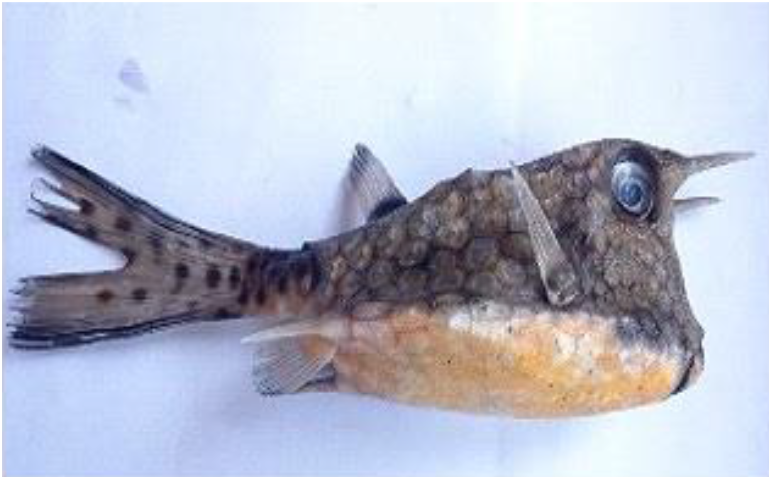
Lateral View.

**Fig 15.0.**
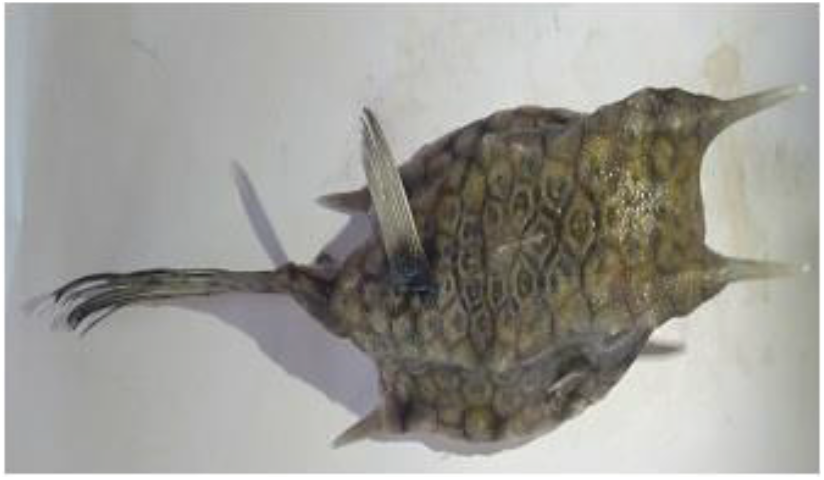
Dorsal View.

**Fig 16.0.**
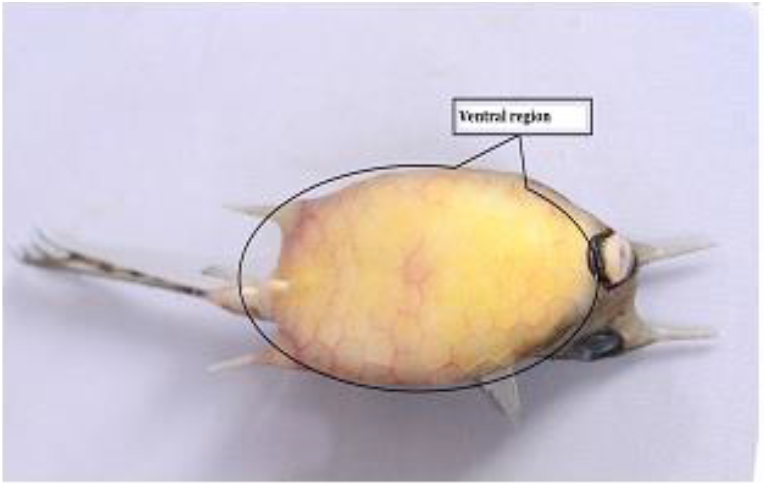
Ventral View.

**Fig 17.0.**
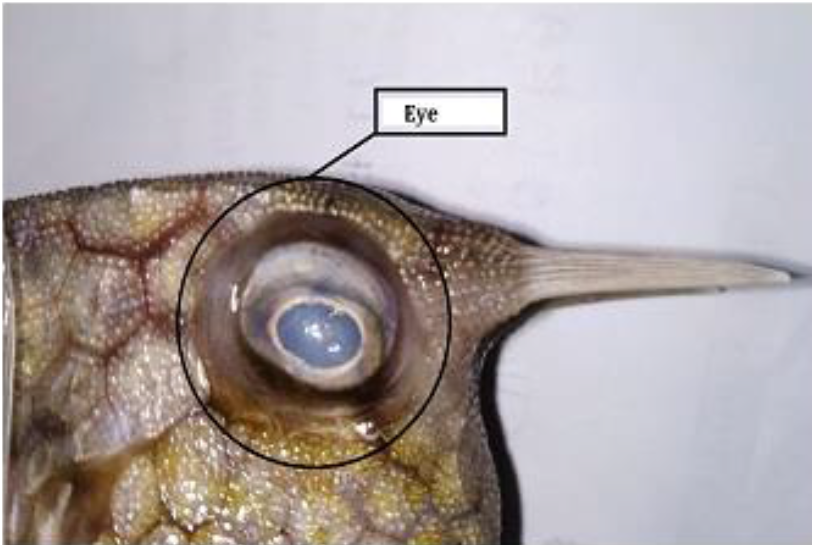
Orbit.

**Fig 18.0.**
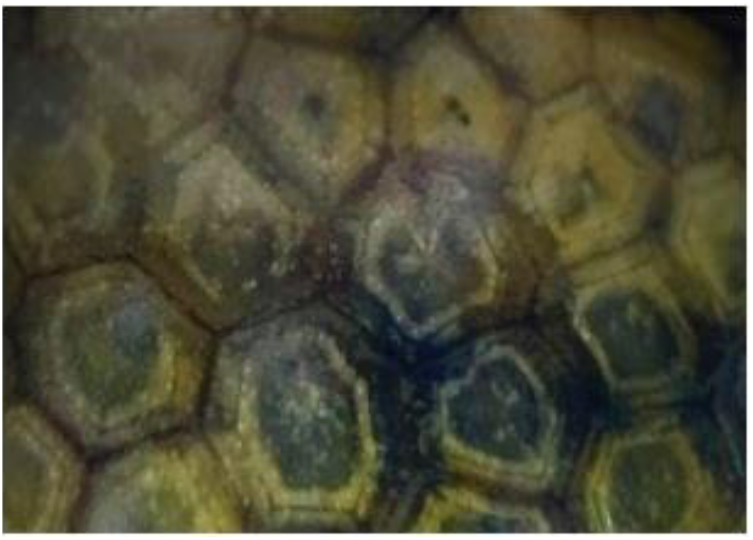
Plate like Hexagonal Scales.

### Second specimen WCI19

Second specimen of *Lactoria cornuta* collected from Majali beach as dead but in fresh and intact condition. All morphometric and meristic studies conducted on WCI18 repeated on this specimen as well (WCI19). The morphometric and meristic data depicted in Table 1.0. and Table 2.0 respectively. In this specimen, the length of right and left anterior as well as posterior dorso-lateral horn lengths found to be equal (Fig.19.0. & 20)

**Fig 19.0.**
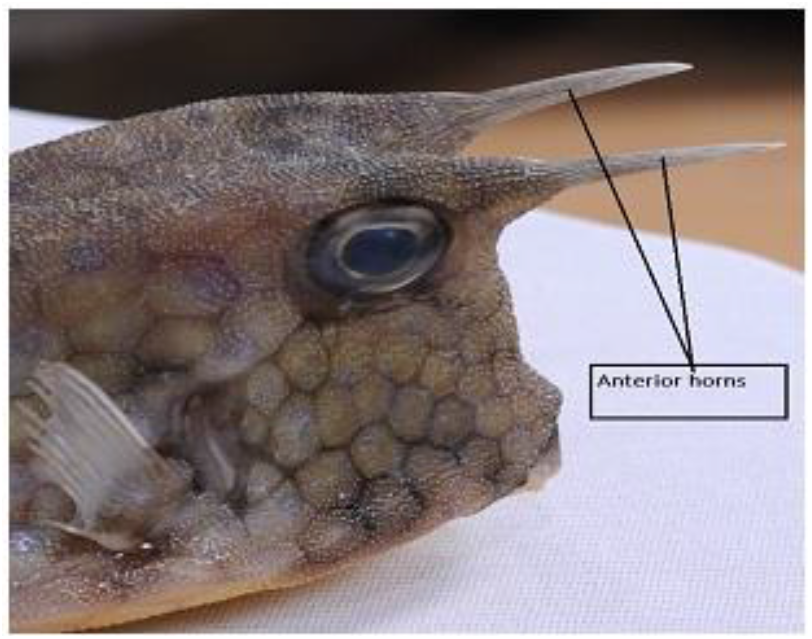
Anterior Horn.

**Fig 20.0.**
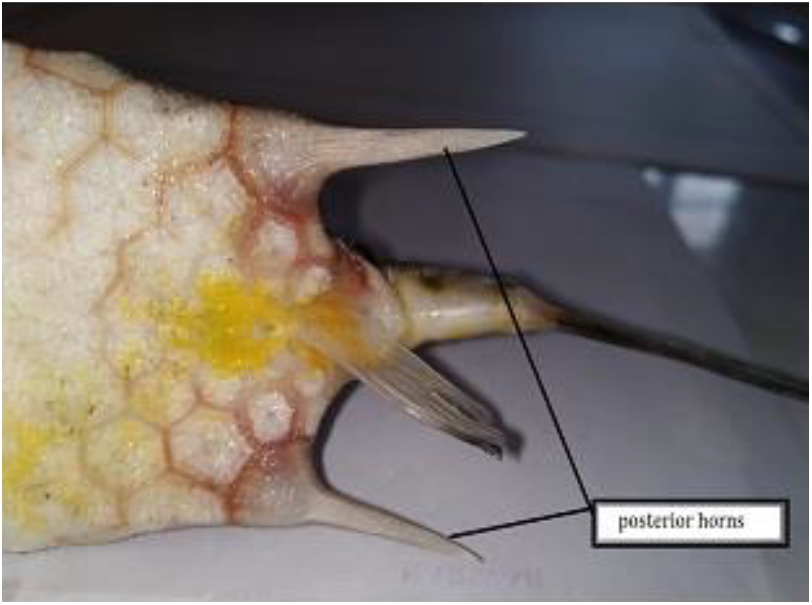
Posterior Hom.

**Fig 21.0.**
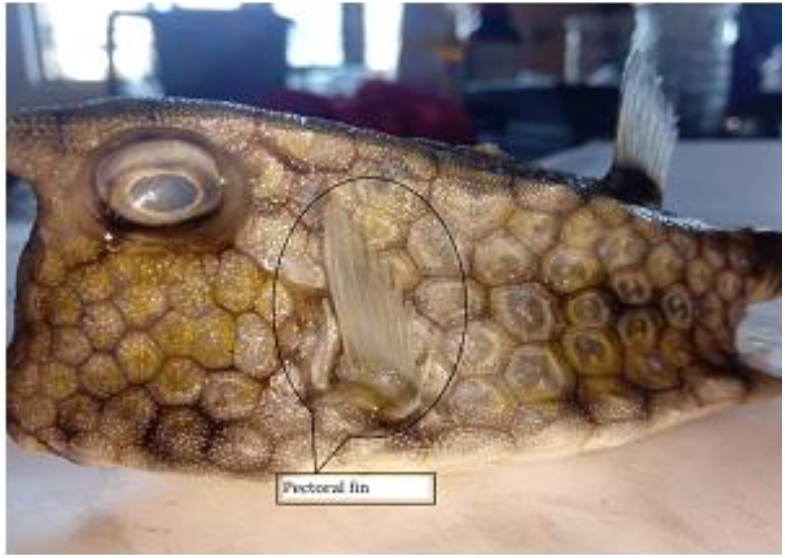
Pectoral Fin.

**Fig 22.0.**
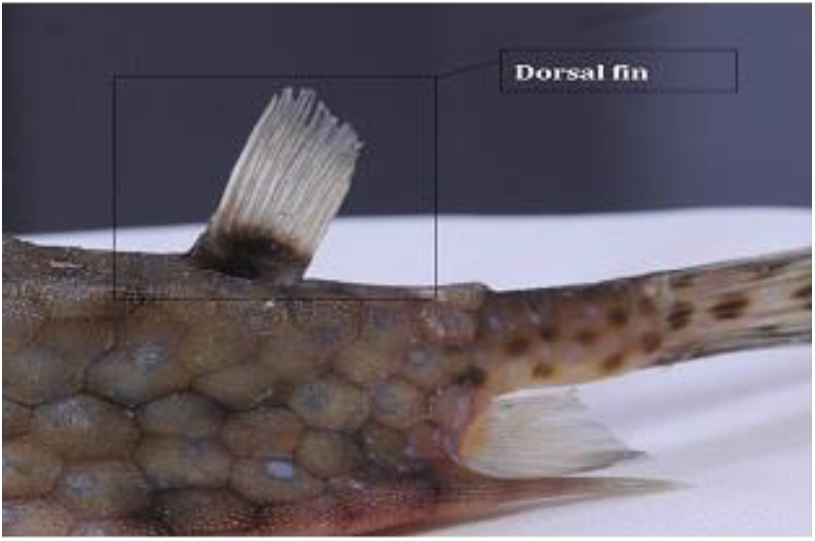
Dorsal Fin.

### Comparative study with the specimens collected from Bay of Bengal

As many as 10 standard morphometric rations calculated not only for our specimens (WCI18 & WCI19) but also for the specimens collected from the Bay of Bengal by other workers. Our aim was to carry out some comparative studies among the specimens collected from the east and west coast of India, it became inevitable for us to calculate the ratios even for their morphometries, as those authors haven’t provided ratios in their publications (Khan, 2013; Behera, 2017). The comparative data of ten important rations given in Table 3.0.

**Table 3.** Comparison of Standard Morphometric Ratios of present study with the specimen collected from Bay of Bengal

| Sl.No. | Measurements | Standard Morphometric Ratios |  |  |  |
| --- | --- | --- | --- | --- | --- |
|  |  | Khan *et.al.*<br>2013 | Behera *et. al.*<br>2017 | Present Study |  |
|  |  |  |  | WCI2018 | WCI2019 |
| 1 | Total Length (TL) :Standard Length (SL) | 0.93 | 1.16 | 1.59 | 1.5 |
| 2 | Total Length (TL) :Head Length (LC) | 6.09 | 5.01 | 6.3 | 5.18 |
| 3 | Standard Length (SL) :Head Length (LC) | 3.95 | 4.38 | 3.95 | 3.87 |
| 4 | Head Length (LC) : Pre Orbital Distance (PRO) | 4.2 | 1.31 | 5 | 4.1 |
| 5 | Maximum Body Depth (Hx) : Minimum Body Depth (Hm) | 1.34 | 1.03 | 1.65 | 1.65 |
| 6 | Eye Diameter (OH) :Inter Orbital Distance (IO) | 0.9 | 0.7 | 0.81 | 0.38 |
| 7 | Total Length (TL) :Pre Dorsal Distance (PD) | 2.56 | 2.64 | 2.37 | 1.68 |
| 8 | Standard Length (SL) :Pre pectoral Distance (PPEC) | 3.6 | 4.51 | 1.49 | 3.55 |
| 9 | Total Length (TL) : Pre Anal Distance (PA) | 2.13 | 2.28 | 2.17 | 2.08 |
| 10 | Standard Length (SL) :Pre Anal Distance (PA) | 1.38 | 1.95 | 1.36 | 1.39 |

In comparative studies, majority of the growth parameters do not show any significant deviations from the specimens collected from the east coast of India (Khan, 2013; Behera, 2017). However, some parameters such as ratios of LC/PRO, SL/PPEC and OH/IO conspicuously differ among the species, the details are provided in Table 3.0.

## DISCUSSION

The longhorn cowfish, *Lactoria cornuta*, is widely distributed in certain pockets of the Indo-Pacific oceans Viz. Red sea, east Africa, Marquesas and Tuamotu Islands, South Korea, Ryukyu Islands (southern Japan), south of Australia and Lord Howe Islands (Goren and Dor, 1994; Van der Elst 1993; Lieske and Myers 1994). Preferred habitats of cowfishes include coral reefs, reef plain habitat, but occasionally they have also been reported from sandy beaches. This species was also reported from the coasts of Indian, it was first reported from Lakshadweep island and later from the coast of Goa as well as from the Bay of Bengal (Vijay Anand & Pillai, 2003; Hegde et al. 2013; Khan, 2013; Behera, 2017).

Apart from the Lakshadweep islands, the longhorn cowfish was only been reported from the single location in the west coast of India (Hegde et al. 2013). In this paper we are reporting the collection of this fish for the first from the coast of Karnataka, west coast of India. We have collected two specimens in two consecutive years, the first specimen was collected in 2018 from Karwar and the second specimen in 2019 in Majali beach (10 Km away from Karwar). Both specimens examined thoroughly and all necessary morphological measurements were taken by following standard methods (Hutchins 1984: Randall et al., 1990; Allen and Erdmann, 2012). This study finds no significant differences in the specimen studied and those from the Bay of Bengal (Khan et al. 2013; Behera et al. 2017). Unfortunately, the specimens collected from the Lakshadweep island and from the coast of Goa were just been documented but never been examined in detail.

The diagnostic characteristics of the specimens collected are consistent with those of the Ostraciid family. The specimen has a typical conical shaped body and covered with a hexagonal plaque like scales, which are fused into box like carapace. Two pairs of anterior and posterior dorsal horns offer enormous survival advantages to escape from enemies, additionally predators find it difficult to swallow due to the presence of horns.(Fig.11 & 19). Other peculiarities included the sub-terminal mouth bound by prominent lips (Fig.1 and 13), gills without operculum. Along with the prominent caudal fin (Fig. 23), these fish also have pectoral, dorsal and anal fins. Range extension of this fish from the selected pockets of the east and west coast of India may either be due to migration or because of other forces such as cyclones and tornados. Given its morphology and body design, migration appears most unlikely to explain for the range extension as such design can only support the lethargic swimming. Several natural forces are known to aid animals in their range extension and in this species, it appears that the super cyclones might have played a role in its range extension.

**Fig 23.0.**
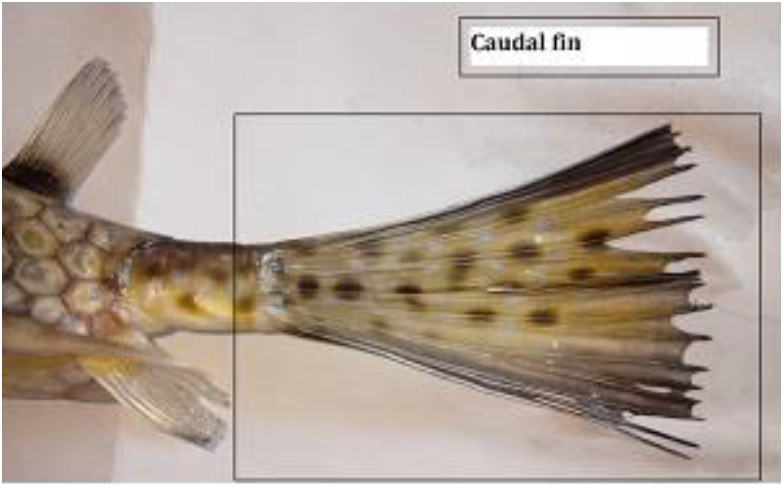
Caudal Fin.

**Fig 24.0.**
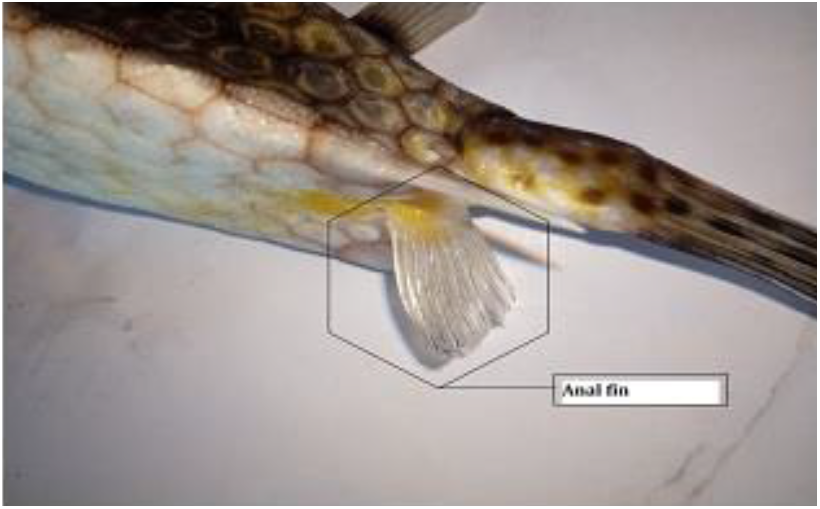
Anal Fin.

**Fig 26.**
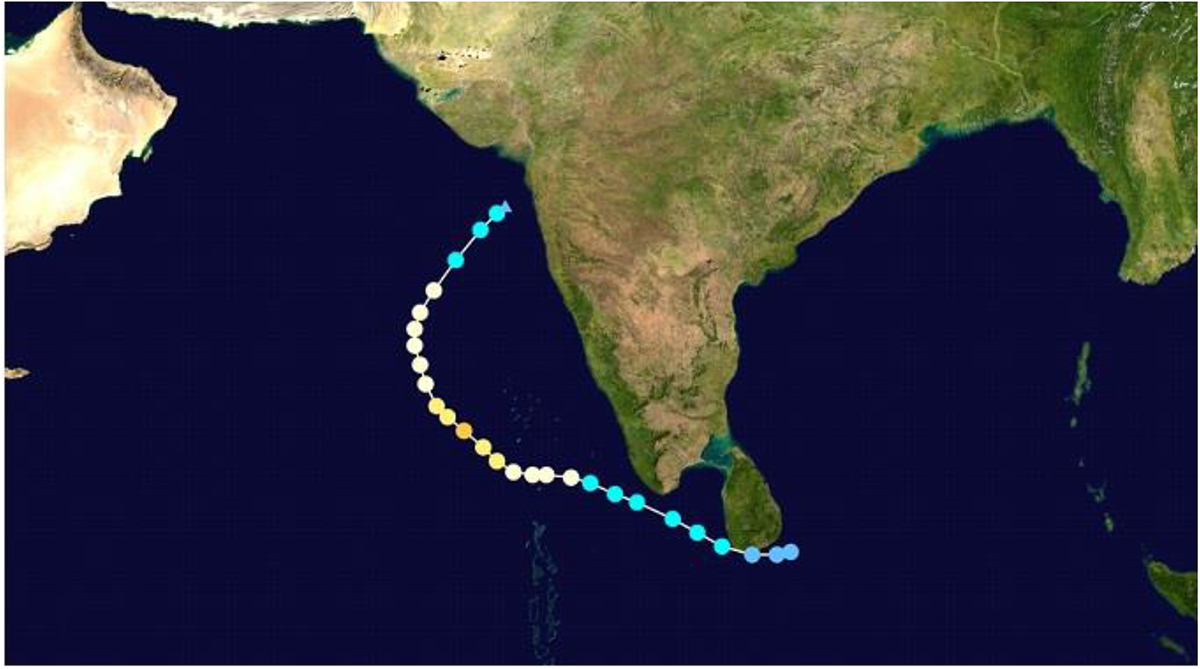
Possible path of drifting of Longhorn Cowfish. (Track created by using http://ports.com)

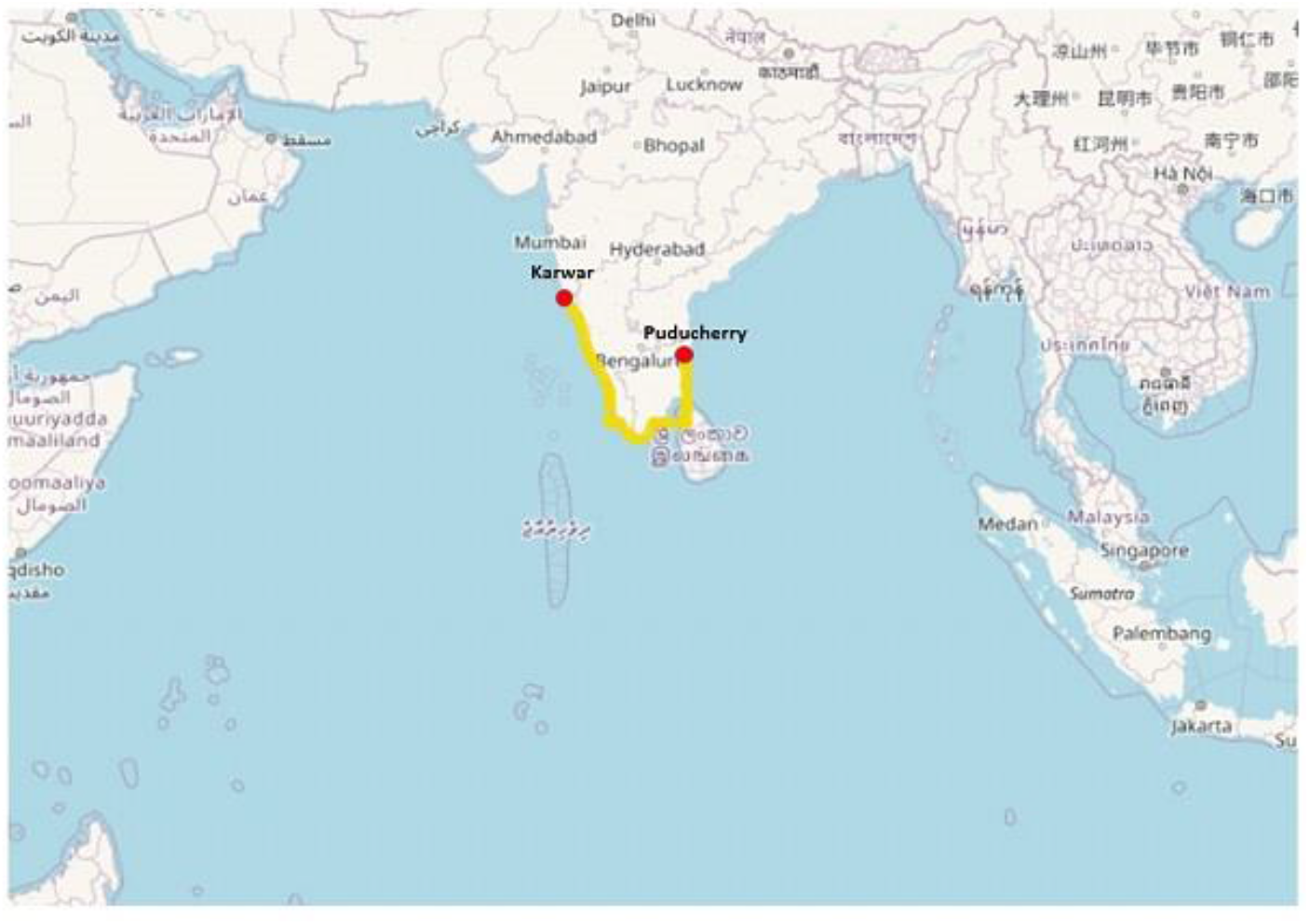

Super Cyclone Ockhi hitting the western coast coincides with the collection of the first specimen along the coast of Karnataka. Therefore, we suggest the role of this super cyclone in the extension of the range of longhorn cowfish from the Bay of Bengal to the west coast of India. Importance of the cyclones in determining the richness, distribution and behaviour of species already been exhaustively discussed in erstwhile studies (Harmelin-Vivien and Mireille, 1994: Gong et al. 2010; Fiedler, et al. 2013; Fandel et al. 2020; Booth and Thomas, 2021; Harley et al. 2006; Easterling et al. 2000; Easterling, 2000). Based on available scientific contributions, we would like to conclude that the cyclones (Track of the most of the cyclones is depicted in Figure 25) must have played a role in determining the range extension of this species.

A year later, another specimen was recovered from a site very close to the first, indicating the successful establishment of *Lacturia cornuta* in its new environment. Relative morphometric and meristic studies of our specimens are consistent with previous studies reported in the Bay of Bengal.

## Acknowledgement

We are grateful to the Department of Collegiate Education, Bangalore, and Government of Karnataka for providing necessary infrastructure for the present study. We have not received any financial assistance for the present studies.

